# Rigid Scaffolds are Promising for Designing Macrocyclic Kinase Inhibitors

**DOI:** 10.1101/2023.03.17.533119

**Authors:** Zheng Zhao, Philip E. Bourne

## Abstract

Macrocyclic kinase inhibitors (MKIs) are gaining attention due to their favorable selectivity and potential to overcome drug resistance, yet they remain challenging to design because of their novel structures. To facilitate the design and discovery of MKIs, we investigate MKI rational design starting from initial acyclic compounds by performing microsecond-scale atomistic simulations for multiple MKIs, constructing an MKI database, and analyzing MKIs using hierarchical cluster analysis. Our studies demonstrate that the binding modes of MKIs are like that of their corresponding acyclic counterparts against the same kinase targets. Importantly, within the respective binding sites, the MKI scaffolds retain the same conformations as their corresponding acyclic counterparts, demonstrating the rigidity of scaffolds before and after molecular cyclization. The MKI database includes 641 nanomole-level MKIs from 56 human kinases elucidating the features of rigid scaffolds, and the tendency of core structures among MKIs. Collectively these results and resources can facilitate MKI development.

## 1. Introduction

The human kinome represents one of the largest gene families, consisting of over 500 protein kinases that regulate almost all aspects of cellular function.^[1, 2]^ Thus, alterations of gene expression, dysregulation of signaling pathways, or gene mutation of kinases can cause a wide variety of cancers and other diseases.^[3]^ Therefore, kinases have been considered primary drug targets.^[4]^ Kinases typically share a conserved catalytic domain that contains a highly similar ATP binding site.^[5]^ As such, it is a daunting challenge to achieve the desired selectivity, where inhibitors bind to the desired kinase but not to the others.^[6, 7]^ Nevertheless, over the last 30 years, a great variety of kinase-targeted inhibitors or degraders have been successfully developed,^[8]^ such as type I/II inhibitors, allosteric inhibitors, covalent inhibitors, macrocyclic inhibitors, PROTAC degraders, and molecular glues.^[9]^ To date, more than 70 small-molecule kinase inhibitors have been approved by the FDA since the first drug Imatinib was approved by the FDA in 2001.^[10]^ Clinically, these drugs have substantially alleviated patients’ anguish and prolonged their lives.^[11, 12]^ However, unexpected side effects and acquired drug resistance also mean that designing more innovative, efficient kinase drugs is warranted.^[2, 11, 13]^

Macrocyclic (at least 12-membered ring) kinase inhibitors (MKIs) have attracted more attention as aids in developing innovative, efficient kinase inhibitors because of their unique cyclic frames and potential to overcome drug resistance.^[8, 14]^ So far, only a few macromolecular MKIs have been published in peer-reviewed journals^[15, 16]^ (namely, Rapamycin and its derivatives binding to the FKBP-Rapamycin binding domain of mTOR kinase,^[17]^ and DNA-template macrocyclic Src inhibitors consisting of peptides^[18]^, **Figure 1**). Here we concentrate on small-molecule MKIs that bind into the ATP-binding pockets. Hitherto, nine small-molecule MKIs have been in clinical trials (www.clinicaltrials.gov), including two FDA-approved drugs Lorlatinib and Pacritinib (**Figure 2**).^[19, 20]^

**Figure 1.**
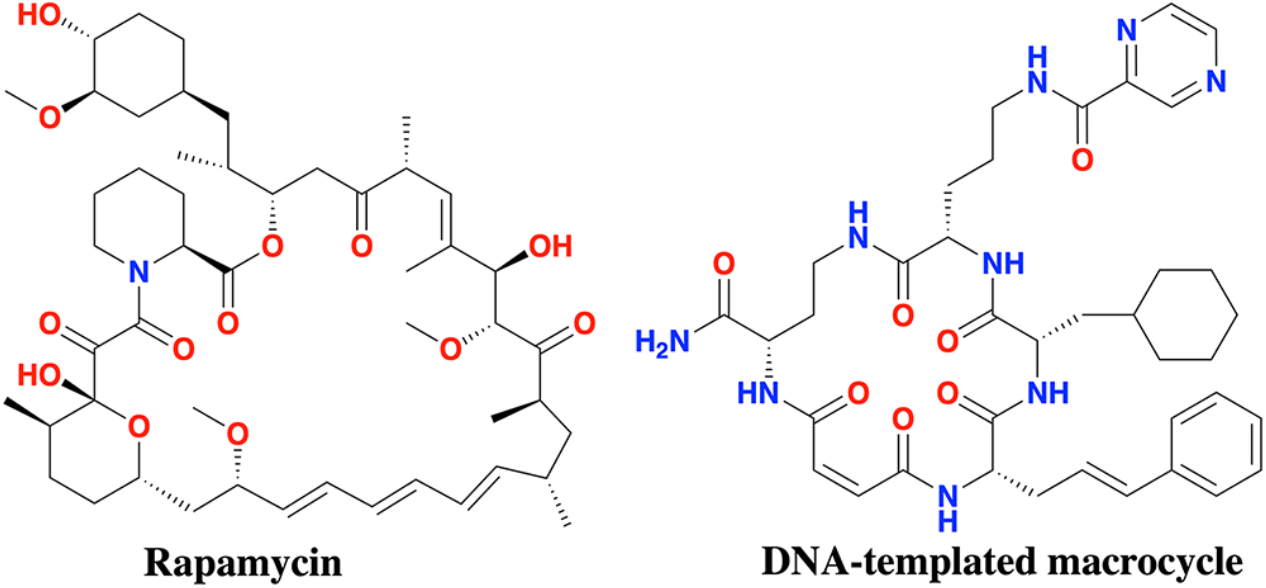
Representative macromolecular MKIs.

**Figure 2.**
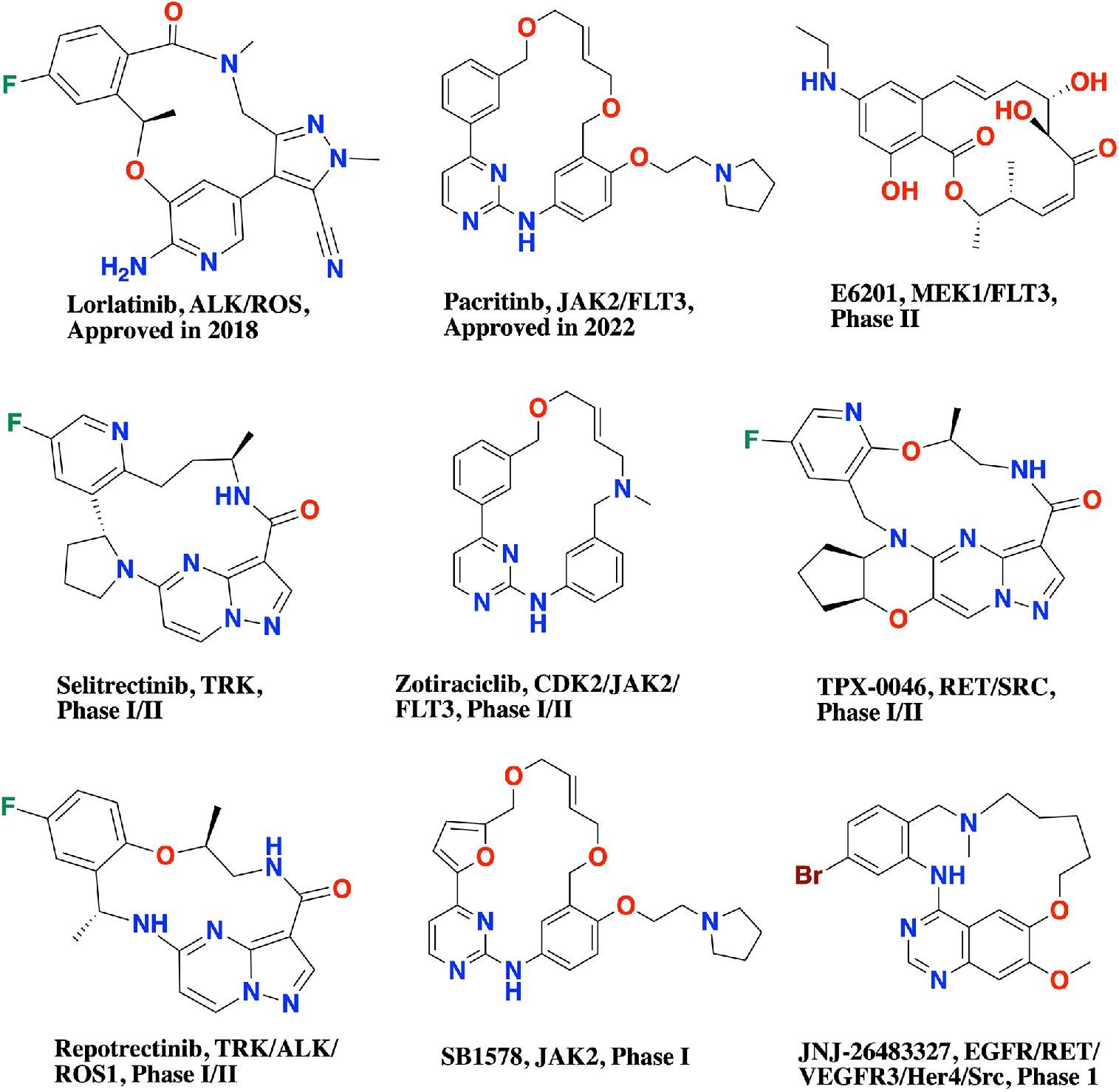
Small-molecule MKIs approved by the FDA or in clinical trials.

Lorlatinib, approved in November 2018, is the first third-generation ALK-targeted drug to overcome multiple recalcitrant resistance mutations, such as G1202R, during the treatment of ALK-positive non-small cell lung cancer (NSCLC) after a year or two of first-or second-generation ALK drug therapy (i.e., Crizotinib, Ceritinib, Alectinib, and Brigatinib).^[20–22]^ Pacritinib, approved in February 2022, is a JAK2/FLT3 inhibitor for the treatment of high-risk myelofibrosis with severe thrombocytopenia.^[19, 23]^ Except for E6201 (**Figure 2**),^[24]^ which was inspired by a natural product called resorcylic acid lactone f152A1,^[25]^ all other eight macrocyclic inhibitors (**Figure 2**) were rationally designed starting from generic acyclic active compounds (called “counterparts”).^[22, 23, 26]^ For example, Lorlatinib was designed based on the acyclic molecule, Crizotinib (**Figure 3**).^[27]^ Pacritinib was designed based on an acyclic, multi-targeted kinase inhibitor, Compound-1 (**Figure 3**). Inhibitor BI-4020 is a fourth-generation EGFR-targeted inhibitor designed to overcome drug resistance acquired from EGFRdel19/T790M/C797M mutations.^[28]^ BI-4020 was designed based on an acyclic, broad kinase inhibitor (here called Ligand-1, **Figure 3**). These successful examples demonstrate that designing MKIs starting from acyclic structures is an effective pathway.^[29, 30]^ However, current *drug-like* chemical space ( at least 10^60^ compounds) is too large and diverse to screen for potent acyclic counterparts that can be used as the starting point in designing MKIs.^[31]^ Narrowing chemical space and determining which acyclic structures are promising for macrocyclic kinase drug design is prerequisite.

**Figure 3.**
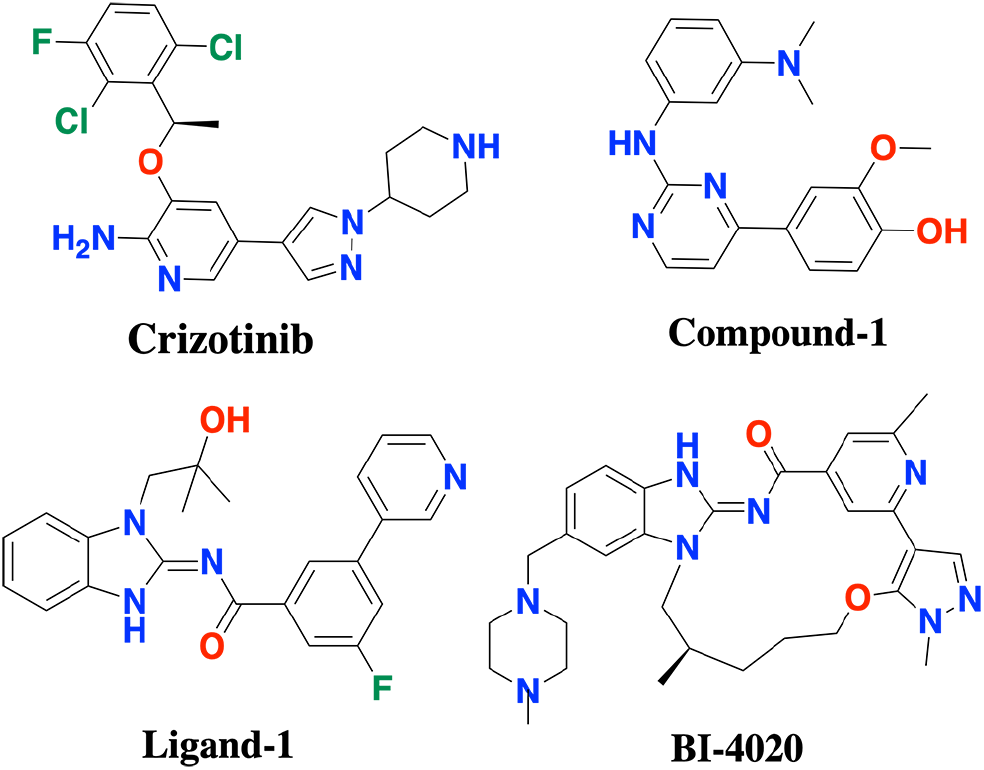
Chemical structures of three acyclic counterpart compounds and MKI BI-4020.

To this end, we sought to explore the molecular characteristics and binding modes of MKIs. We performed microsecond-scale atomistic simulations for three pairwise systems (i.e., Lorlatinib and Crizotinib,^[22]^ Pacritinib and Compound-1,^[23]^ and BI-4020 and Ligand-1,^[32]^ **Figure 3**) to identify the different binding characteristics between MKIs and their corresponding acyclic counterparts within the kinase binding pockets. Based on these simulations, a systematic analysis of the binding characteristics of MKIs before and after their cyclization was performed using a function-site interaction fingerprint (Fs-IFP) method.^[33, 34]^ Subsequently, we manually constructed an MKI database from the published MKI literature. Thus far, a total of 641 nanomolar MKIs, covering 56 human kinases, have been curated. The data can be accessed and downloaded freely from (https://zhengzhster.github.io/MKIs). Harnessing the MKI database, we got an overview of the properties of MKIs, paying particular attention to the characteristics of MKI scaffolds. In addition, the core structures of MKIs that typically interact with the hinge regions are discussed, and design strategies are proposed. Together the binding modes, database, and design strategies for MKIs provide a resource for advancing MKIs’ design and discovery.

## 2. Results

### 2.1. Binding Modes of MKIs and Their Corresponding Acyclic Counterparts

We first analyzed the binding modes of MKIs and their corresponding acyclic counterparts within the kinase binding sites using microsecond-scale all-atom MD simulation (See the **Method** section).^[33, 35]^

#### Binding modes of Crizotinib and Lorlatinib

The binding modes of the two ALK inhibitors are illustrated in **Figure 4a-b**. The common aminopyridine cores of Crizotinib and Lorlatinib form stable interactions with residues E1197, L1198, M1199, and L1256 (the probabilities of interactions > 0.8, **Figure 4b**). Of these, E1197, L1198, and M1199 are at the hinge. L1256 is beneath the aminopyridine core. Thus, the aminopyridine scaffold, like the adenine ring of ATP, binds into the ATP-binding site and forms at least one hydrogen-bond interaction with the hinge.^[22, 27]^ Linking to the aminopyridine core, the common fluorophenyl groups of Crizotinib and Lorlatinib lies between the roof (β3) and the DFG peptide with similar interactions with V1130, K1150, N1256, and D1270. Another common pyrazole moiety, connecting the aminopyridine core, is located between L1122 and G1202. However, in Crizotinib, the piperidine-substituted pyrazole moiety is exposed to the solvent, while interacting with G1123, A1200, and S1206 in the front-pocket area (**Figure 4a-b**). By contrast, in Lorlatinib, the pyrazole was optimized forming 2-methylpyrazole-3-carbonitrile, in which the nitrile group interacts with R1120 and E1132.^[22]^ In summary, the core scaffolds of Crizotinib and Lorlatinib present similar binding modes with differences in the substructures,

**Figure 4.**
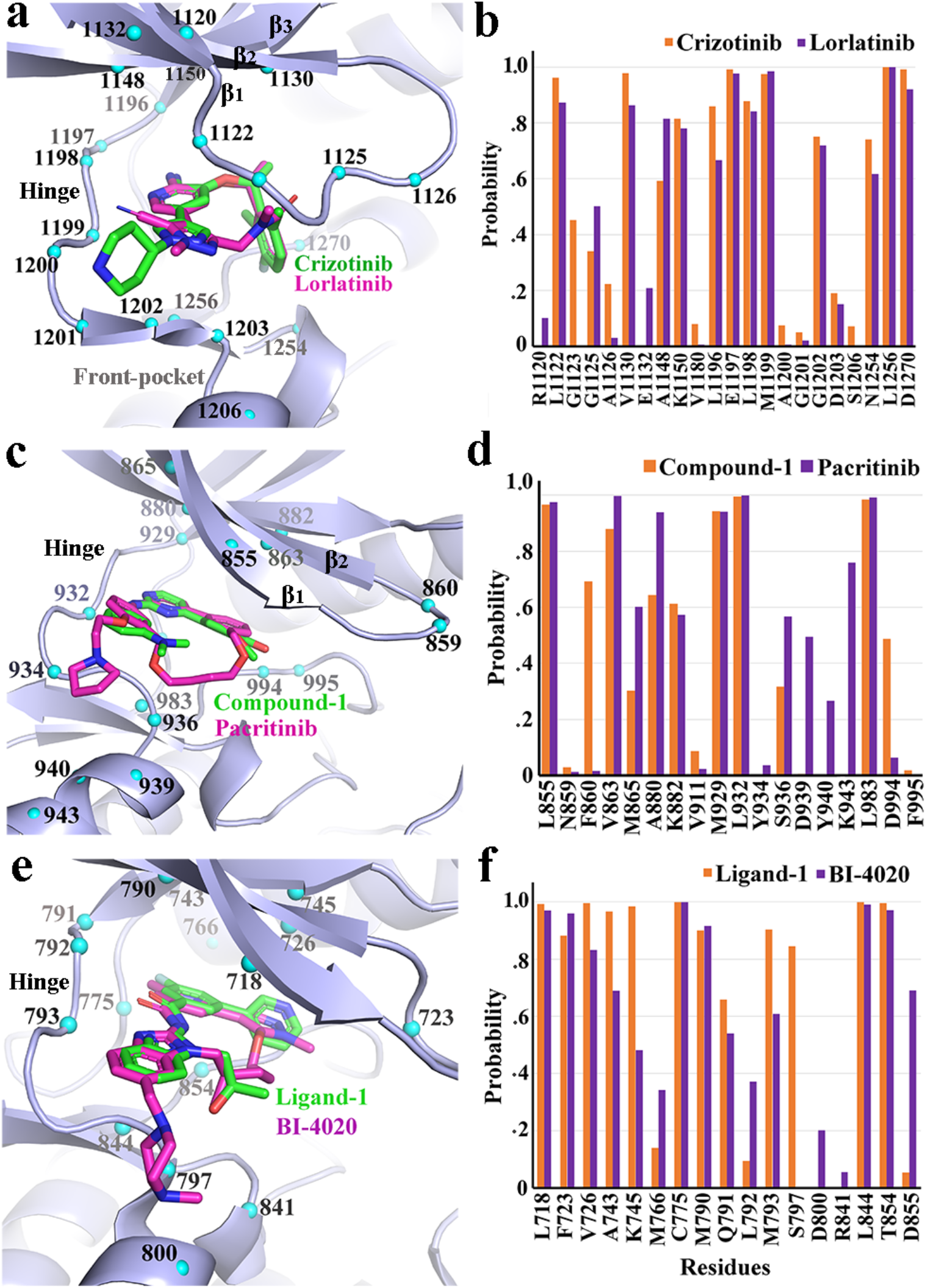
(a-f) The binding modes of each pairwise system (MKIs and their corresponding acyclic counterparts; PDB templates: 5aa9, 7ree, 7kxz for a, c, and e, respectively).

#### Binding Modes of Compound-1 and Pacritinib

Pacritinib, a JAK2-targeted drug, was designed based upon an acyclic counterpart, Compound-1, which was then optimized by cyclization and R-group modification.^31^ Thus, Pacritinib and Compound-1 share the same scaffold, which is composed of an aminopyrimidine group coplanar with two phenyl groups (**Figure 3, 4c**). In Compound-1, the aminopyrimidine core structure, located at the adenine binding area, has stable hydrogen-bond interactions with residues M929 and L932 at the hinge (probability of interaction > 0.9, **Figure 4c-d**). The 2-methoxyphenol fragment, connecting to the aminopyrimidine core, forms stable hydrophobic interactions with residues V863 at the β2, and L983 at the β6 strand. The terminal dimethylphenylamine moiety makes stable hydrophobic interactions with residue L855 at the β1 (probability of interaction > 0.9, **Figure 4d**). Compared to Compound-1, the two open ends between the two phenyl groups in Pacritinib are bridged. After cyclization, the binding patterns between the Pacritinib scaffold and residues L855, V863, M929, L932, and L983 are similar to that of Compound-1 (**Figure 4d**). Apart from the common scaffolds Pacritinib was further optimized through a terminal pyrrolidine displacement on the 4-aminophenol fragment. The pyrrolidine extends into the solvent front-pocket interacting with residues D939, Y940, and K943, accounting for the improved selectivity against JAK2.^[19]^

#### Binding Modes of Ligand-1 and BI-4020

BI-4020 is a potential fourth-generation EGFR inhibitor that overcomes del19/T790M/C797S mutation-induced drug resistance.^[32]^ Based on the scaffold of Ligand-1, BI-4020 was rationally designed. The common scaffold of the two share a similar binding mode of Type-I kinase inhibitors (**Figure 4e**). Specifically, the common aminobenzimidazole core has stable interactions with residues Q791, L792, or M793 at the hinge, residue L718 at β1, and residue L844 at β6 (Probability > 0.9, **Figure 4f**). The “head” groups of both, attached to the aminobenzimidazole group, extend into the kinase hydrophobic sub-pocket, forming key interactions with residues A743, M766, C775, and T854, respectively (**Figure 4f**). In contrast, the terminal piperazine group of BI-4020 reaches the solvent, forming interactions with residue D800 at the front pocket. Moreover, the bridging linker of BI-4020 also provides unique interactions with residues R841 and D855 to achieve selectivity among mutant variants (**Figure 4f**).

Overall, MKIs and their corresponding acyclic counterparts have the same Type-I binding modes, forming the typical hydrogen-bond interactions with the hinge.^[36]^ The same scaffolds of MKIs and their corresponding acyclic counterparts show similar binding patterns, meaning the binding modes of the scaffolds are not affected upon macrocyclization. We investigated the fluctuation of ligands within the binding sites by analyzing the MD trajectories of MKIs and their corresponding acyclic counterparts. This showed that the scaffolds remain more rigid than other fragments of the ligands either before or after macrocyclization (**Figure 5**). Considering the consistency of scaffolds before and after cyclization we further studied the properties of MKIs, to explore promising new scaffolds for developing MKIs.

**Figure 5.**
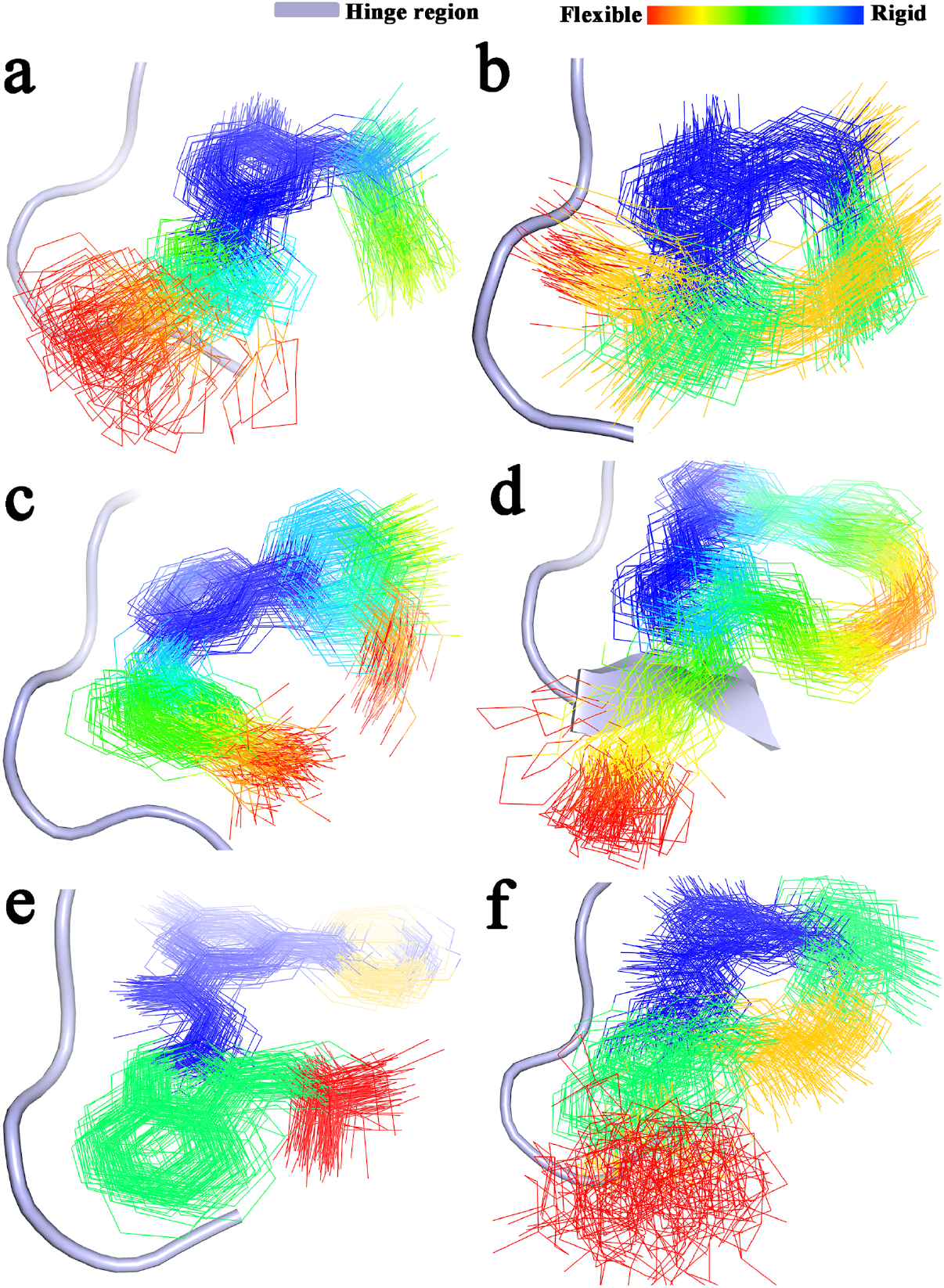
(a-f). The rigidity of Crizotinib, Lorlatinib, Compound-1, Pacritinib, Ligand-1, and BI-4020, respectively, while interacting with the hinge (grey). Rigid scaffolds to flexible fragments are colored from blue to red.

### 2.2. Characteristics of MKIs and Scaffolds

The consistent rigid properties of the scaffolds before and after the aforementioned cyclization inspired us to establish the characteristics of such scaffolds so they may be used to screen MKIs from across acyclic chemical space. We collected all of the released MKIs as found in the literature constructing an MKI database containing 641 MKIs and 56 human kinases with nanomolar affinity (**Figure 6a**, **Table S1-S2**). The 56 kinase targets were belonged to different kinase groups (TK, CMGC, CAMK, AGC, STE, TKL, Other, Lipid, and Atypical) excluding the CK1 group; 20 out of 56 belong to the TK group. The two kinase targets (ALK and JAK2) with the approved macrocyclic kinase drugs, Lorlatinib and Pacritinib. belong to the TK group. We obtained the scaffold of every MKI using the tool strip-it.^[37]^ After deleting the redundant scaffolds, a total of 95 unique scaffolds were obtained (**Table S3**), and then clustered using an “average linkage” hierarchical clustering algorithm.^[38]^ Ten scaffold clusters were identified using a dissimilarity distance > 0.8 as the threshold (**Figure 6b, Table S4**). The 10 clusters provide an available conformational space for designing MKIs with diverse properties. The details reveal the important conformations and properties.

**Figure 6.**
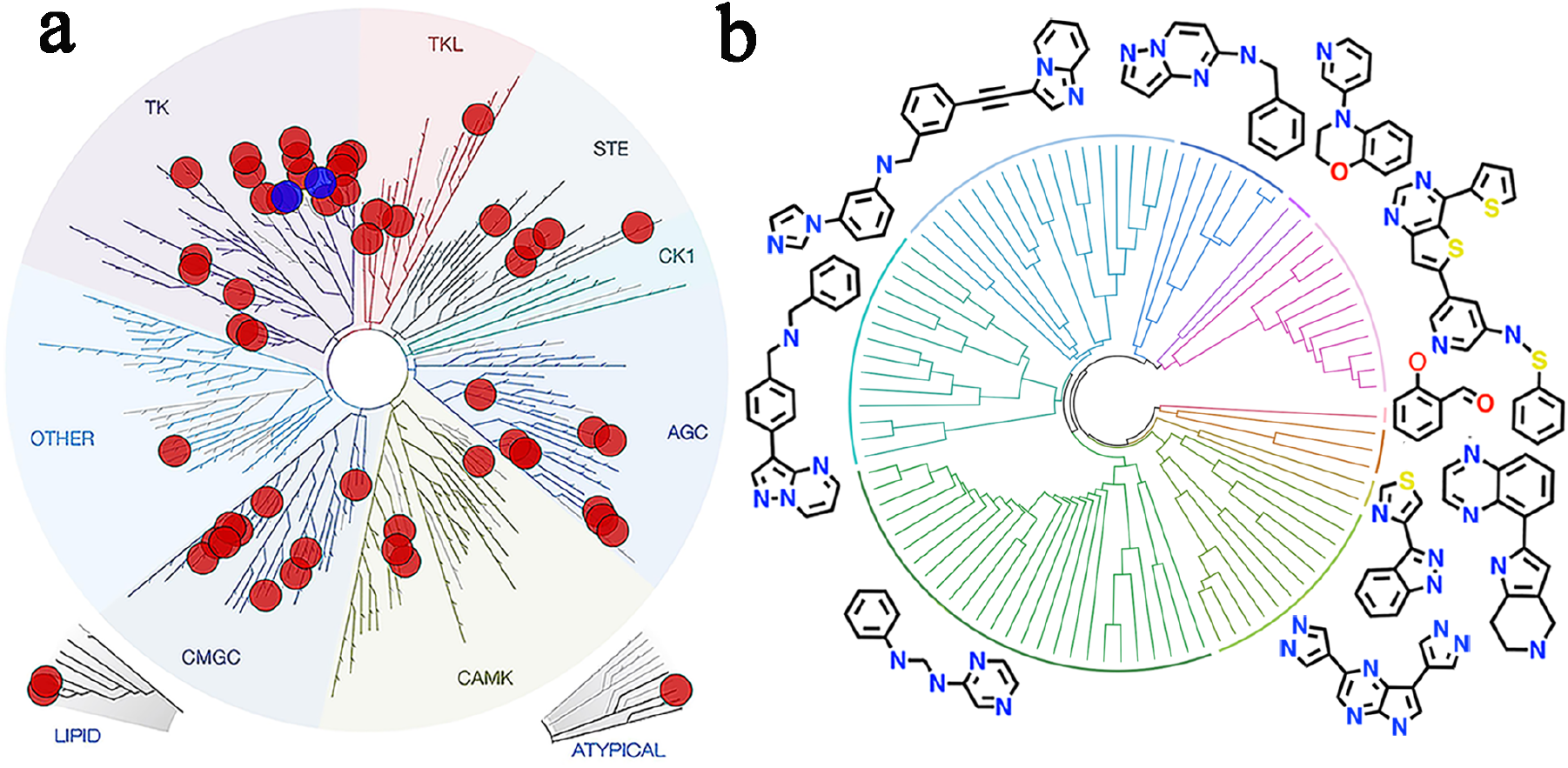
(a) Distribution of kinase targets with released MKIs across the human kinome (red). Two kinases with approved macrocyclic kinase drugs are highlighted in blue. (b) Distributions of ten clustered scaffolds with one compound representative per cluster.

For every scaffold and MKI, molecular weight (MW), number of H-bond donors (HbD), number of H-bond acceptors (HbA), logP, number of aromatic rings (AR), and molecular flexibility (MF) have been analyzed (**Figure 7, Table S5-6)**. For MKIs, the MW ranges from 296 to 720 g. mol^-1^, the number of HbDs from 0-6, the number of HbAs from 3-10, the logP has a range of [-6.01, 8.74], the number of ARs from 1-5, and the MF from 3.62 to 15.85. The averages for MWs, HbDs, HbAs, logPs, ARs, and MFs of MKIs are 457.4, 2.3, 6.3, 1.9, 3.0, and 7.7, respectively. According to Lipinski’s rule of five (Ro5, **Table S7**), 99.4% of MKIs have HbD values of ≤ 5, 100% of MKIs have HbA values of ≤10, and 96.1% of MKIs have logP values of ≤ 5. Notably, only 74.5% of MKIs have an MW of ≤ 500, which is in agreement with the trend in MW increase for FDA-approved drugs over the last 20 years.^[39]^

**Figure 7.**
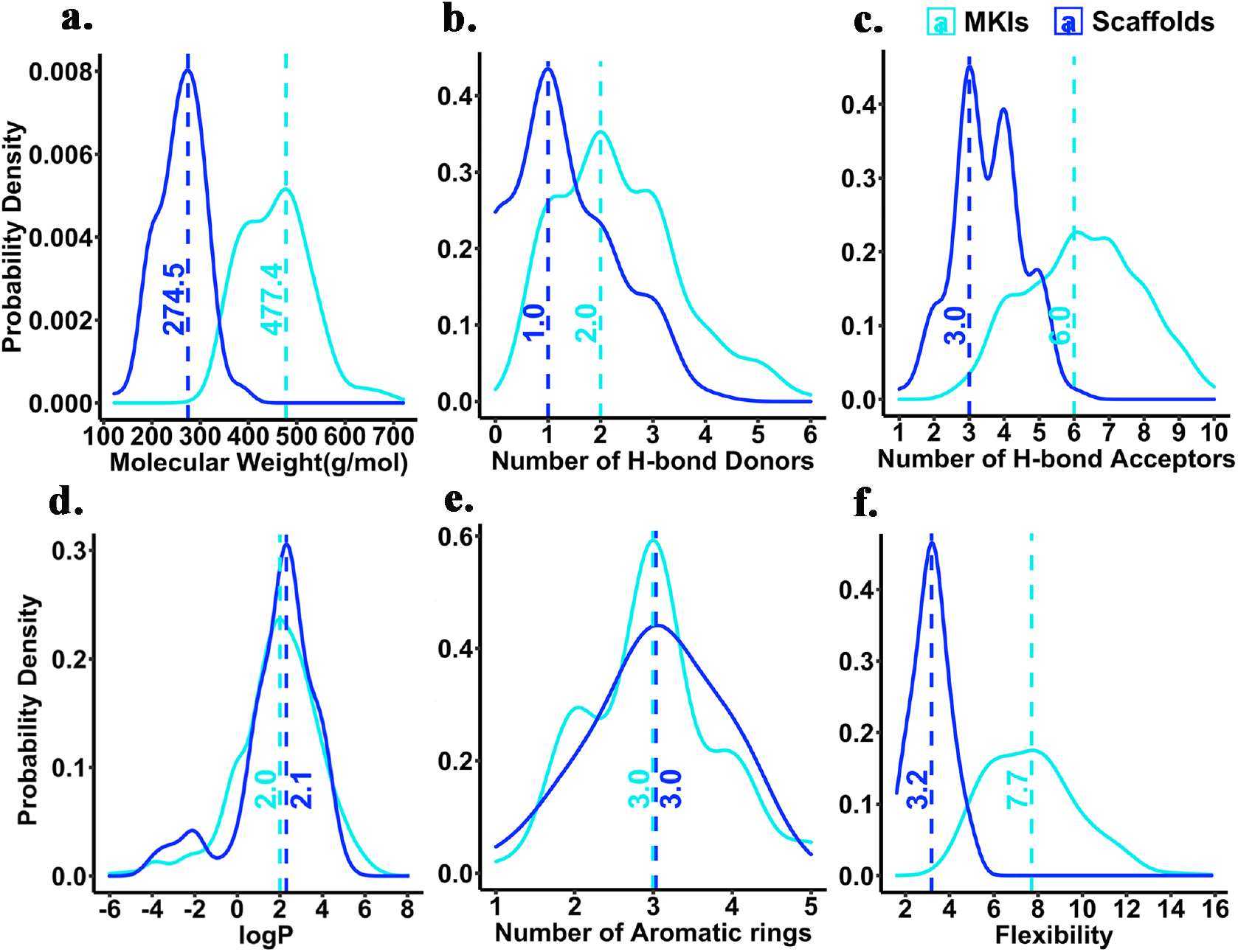
(a-f) Properties of MKIs and the corresponding scaffolds: molecular weight, number of H-bond donors, number of H-bond acceptors, logP, number of aromatic rings, and molecular flexibility, respectively. The ordinate is a probability density. The dashed line indicates the maximum probability density and the corresponding abscissa.

The maximum probability densities (MPDs) for MW, number of HbD, and number of HbA of scaffolds are 274.5, 1.0, and 3.0, respectively, smaller than those for MKIs (477.4, 3.0, and 6.0) (**Figure 7a-c**). Not surprising since the ring_with_link scaffolds are obtained by excising the bridging linkers and deleting the R-groups of MKIs.^[37]^ The logPs of scaffolds and MKIs have similar MPDs (2.1 and 2.0), which mean that both have similar hydrophobicity profiles.^[40]^ From this point of view, it may be necessary to consider hydrophobicity when designing MKIs starting with the choice of scaffold. Likewise, the number of ARs for scaffolds and MKIs are similar with MPDs of 3.0 (**Figure 7e**), suggesting that ARs are part of scaffolds but no ARs are in bridging linkers for MKIs. The scaffolds have much low flexibility (3.2) than MKIs (7.7) (**Figure 7f**). Not surprising since trimming the R-groups of MKIs to obtain scaffolds, the MFs of scaffolds would decrease. Scaffolds consist of a few rigid ARs (**Figure 7e**) consistent with the aforementioned rigid binding modes. More specifically, since the number of ARs and MFs of scaffolds have similar MPDs (3.0 vs 3.2), it means scaffolds are mostly composed of 3 ARs directly connected to one another. This suggests that ARs-composed scaffolds with strong rigidity should be considered for rational MKI design. The majority of MKIs obey the Ro5, and when combined with the characteristics of MKIs it suggests that multiple-AR, rigid scaffolds are practicable in developing MKIs.

### 2.3 Strategies to Design MKIs from Core Structures that Interact with the Hinges

To date, all published MKIs are type I or type II kinase inhibitors that occupy the ATP-binding site and form hydrogen bond interactions with the hinge.^[6]^ While the hinge-MKI interaction appears to be conserved, the types and positions of the atoms involved in hydrogen bonding can vary significantly. Accordingly, understanding what chemical structures can serve as the core structures for recognizing and binding into the adenine sub-pocket is essential to choreograph scaffolds for MKI design. Hence, we investigated all MKIs based on differences in core structures interacting with the hinge, establishing 73 different core structures (**Table S8**). The 73 core structures were further categorized into eight groups (**Figure 8**). In the dendrogram, each node represents a core structure, and edges connect nodes that share the same chemical fragments interacting with the hinge (**Figure 8**).

**Figure 8.**
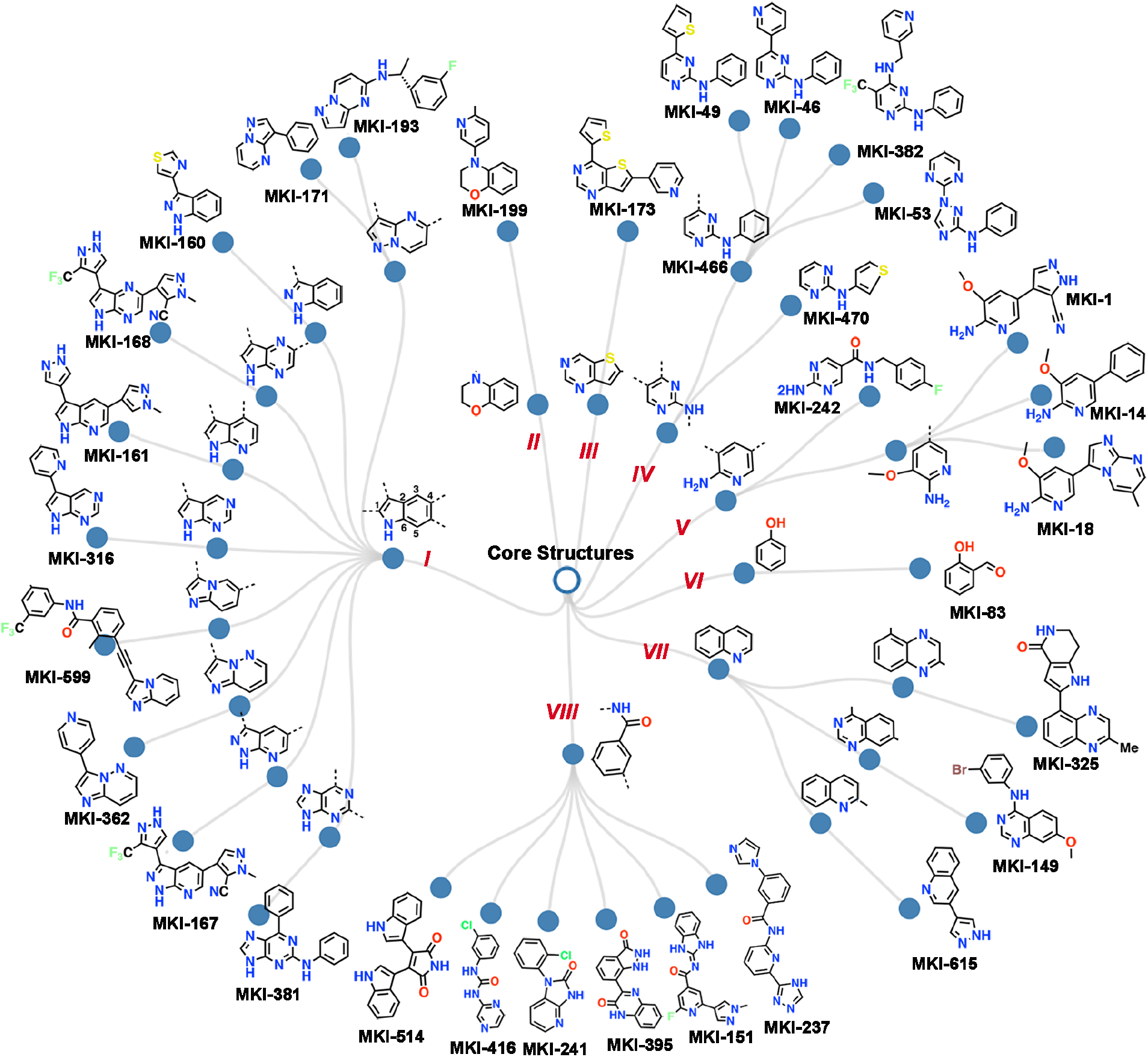
Eight MKI core structure categories. One representative per node (The full list is in **Table S8**).

Specifically, the eight groups are: *I:* indole derivatives; *II:* benzomorpholine derivatives; *III:* thienopyrimidine derivatives; *IV:* 2-amino-pyrimidine derivatives; *V:* 2-amino pyridine derivatives; *VI:* phenol derivatives; *VII:* quinoline derivatives; and *VIII:* benzamide derivatives. On the innermost layer, the nodes represent common fragments interacting with the hinge for every group. The outer-layer nodes show examples of structural derivatives in groups *I-VIII*, respectively. For example, in group V, the innermost layer is 2-amino pyridine, in which the amino and the nitrogen atom of the heterocyclic pyridine provide the hydrogen-bond interactions with the hinge. Moreover, the amino group is located on the side close to the gatekeeper.^[36, 41]^ In the middle layer, the derivatives were first divided into two prongs: the first is a 5-substituted 2-aminopyrimidine derivative (**MKI-242**), and the second branch is 3,5-substituted 2-aminopyridine derivative. Then, based on the 5-substituted difference, the second branch was further derived into three prongs in the outer layer with representatives **MKI-1**, **MKI-14**, and **MKI-18**.

Similarly, the other seven groups illustrate different categories of core structures interacting with the hinge. In the largest group, group *I*, the indole nitrogen atom provides the hydrogen bond interaction with the hinge. Given heterocycles with more nitrogen atoms, such as in the 1, 2, 3, 4, 5, or 6 position, the group diverges into 9 branches: pyrazolo[1,5-a]pyrimidine, 1H-indazole, 5H-pyrrolo[2,3-b]pyrazine, 7-azaindole, 7H-pyrrolo[2,3-d]pyrimidine, imidazo[1,2-a]pyridine, imidazo[1,2-b]pyridazine, 7-azaindazole, and purine, as shown on the middle layer. Notably, they share the same planar ring structures akin to adenine, but the different heterocycles provide opportunities for scaffold hopping for drug design.^[42]^ It is not surprising that adenine-like structures tend to appear in adenine pockets and this suggests that adenine-like core structures are practical MKI scaffolds.

## 3. Conclusions

Here we focus on MKIs, a class of emerging kinase inhibitors with novel molecular scaffolds, that provide potential opportunities for establishing new chemical entities through studying the binding modes of MKIs in kinase pockets, curating a MKI database, and determining the characteristics of MKIs’ scaffolds.

We first explored the binding modes of MKIs against kinase targets and compared them to their corresponding acyclic counterparts by performing three pairs of *μs*-scale MD simulations. Our simulations show that the binding modes of MKIs and their acyclic counterparts are similar. The binding patterns of scaffold fragments before and after cyclization retain higher rigidity and consistency, inspiring us to pay more attention to acyclic compounds, especially, those that can be used for MKI design. To this end, we investigated MKI’s characteristics by creating a MKI database, containing 641 MKIs covering 56 kinases. Based on analysis of the MKI database, we found 95 diverse scaffolds where a large proportion obey the Ro5. Compared to MKIs, the 95 scaffolds have the same logP profiles, which suggests that we should emphasize solubility and lipophilicity when designing MKIs. The scaffolds are generally composed of 3 ARs connected by one bond, which makes for rigid MKIs’ scaffolds. Further, according to typical hinge-ligand interactions of type-I/II kinase inhibitors, we investigated the core structures of MKIs and demonstrated that adenine-like core structures tend to appear in the adenine pockets.

In summary, this work systematically studied MKIs, revealing the promise of rigid scaffolds for designing MKIs and a preference of adenine-like core structures for developing scaffolds. The MKI database used in this study is freely available at https://zhengzhster.github.io/MKIs. The resultant database and understanding of the chemical characteristics can go a long way to reducing the vast chemical space needed for MKI screening.

## 4. Experimental Section

### 4.1. MD Simulations

All-atom MD simulations have been widely used for exploring drug binding mechanisms.^[43]^ Here, the goal is to explore the binding modes of macrocyclic inhibitors before and after cyclization using microsecond-scale MD simulations. To do so, three pairs of MD systems for both the macrocyclic inhibitors and their corresponding counterparts (i.e., the initial acyclic hit compounds for starting to design the MKIs) were prepared. Lorlotinib was rationally designed based on an acyclic drug Crizotinib.^[22]^ Thus, the MD simulations of Crizotinib and Lorlatinib bound to ALK kinase were set up and their starting conformations were taken from Protein Data Bank (PDB)^[44]^ (PDB ids: 2xp2 and 5aa9, respectively). Likewise, the starting conformations of BI-4020 and its corresponding counterpart (named Ligand-1)^[32]^ bound to EGFR kinase were taken from the PDB (PDB ids: 7kxz and 6s9b, respectively). Because the cocrystal structures of the macrocyclic drug Pacritinib and its counterpart (named Compound-1)^[23]^ bound to JAK2 kinase weren’t available in any protein structure database, we docked the two molecules Pacritinib and Compound-1 into JAK2 ATP binding pockets, respectively (PDB id 7ree as the kinase template) using the AutoDock4.2 software.^[45]^ From the docked lists of complexes, the top scoring complexes were selected as the starting conformations of the Pacritinib- and Compound-1-bound JAK2 systems, respectively.

All six kinase-ligand complexes were processed using the VMD software,^[46]^ with missing residues added based to the corresponding kinase sequence. Redundant structures were deleted based on the amino acid sequence of each kinase domain (ALK residues 1116-1392, EGFR residues 712-979, and JAK2 residues 849-1124). Mutations at E746_A750del (representing the del19 mutations^[47]^), T790M, and C797S were set for the EGFR kinase system as BI-4020 is sensitive to the triple mutation while sparing wildtype EGFR.^[32]^ These 6 systems were solvated in a rectangular water box with margins at an 18.0Å buffer distance from any solute atom. The protonation states of all charged amino acids were automatically assigned assuming a PH of 7.0. The counterions (Na^+^ and Cl^-^) were added to reach establish an ion concentration of 0.20 M and electroneutrality. The CHARMM36 all-atom protein force field,^[48]^ CHARMM general force field,^[49]^ and TIP3P water model were used to describe kinases, ligands, and water molecules, respectively. The parameter files and the topology files of all ligand molecules (i.e., Crizotinib, Lorlatinib, Compound-1, Pacritinib, Ligand-1, and BI-4020) were prepared using the online server (https://cgenff.umaryland.edu/).^[50]^

For MD simulations, every system was first optimized using an ACEMD MD protocol: 500 steps of minimization and a 5 × 10 ns restraining MD simulation with gradual reduced restraining force constants (i.e., 10.0, 5.0, 2.5, 1.0 and 0.5 kcal.mol^-1^.Å^-2^, respectively).^[51]^ A 1.2 *μs* MD simulation was carried out (**Figure S1**) and the last 0.8 *μs* equilibrated MD trajectory was used to analyze the ligand-binding details for each system. All MD simulations were performed using the ACEMD software package.^[51]^ During the MD simulations, the integration time step was 4fs, and the SHAKE method was used as the bond constraint.^[52]^ The temperature and the pressure were maintained using a Langevin thermostat at 298.15K and a Berendsen barostat at 1atm, respectively. All trajectories and the ligand fluctuations were analyzed using a Wordom program.^[53]^

### 4.2 Kinase-ligand Interaction-fingerprint Analysis

As stated, all conformations were derived from the last 0.8 μs equilibrated MD trajectories for each system in order to analyze the binding characteristics. Specifically, we encoded every conformation into a one-dimension array as a bit string,^[33]^ representing the kinase-ligand atom-scale interaction details, by using an Fs-IFP method with predefined geometric rules.^[34, 35, 54]^ In the Fs-IFP method, first, all kinase-ligand conformations were aligned using a 3D binding pocket alignment tool SMAP.^[55]^ Second, in the ligand-binding binding site, every residue-ligand interaction was described using 7 kinds of interaction fingerprints (IFPs): van der Waals, aromatic interaction (face-to-face or face-to-edge), hydrogen bond (protein as acceptor or donor), and electrostatic interactions (protein positively charged or negatively charged).^[56]^ Thus, the interactions between every residue within the binding site and the ligand were encoded into a 7-bit substring, such as “1000000”, where “1” indicates the interaction exists and “0” means no interaction detected between the given residue and ligand. The residue-based bit string was encoded for every kinase-ligand conformation. Finally, based on the aligned binding pockets, all Fs-IFPs were aligned for analysis. The IChem software package was used for encoding IFPs.^[56]^

The probability of interaction between the ligand and every residue comprising the binding sites was calculated. From MD simulation trajectories, the probability of interaction of every residue is obtained using the equation 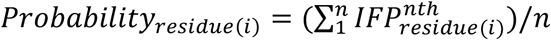, where *i* is the index of amino acids comprising the binding site; *n* is the number of conformations extracted from the corresponding MD trajectory. In conformation *nth*, if a 7-bit substring between the *residue* (*i*) and the ligand is “0000000”, which means no interactions, 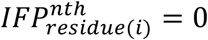, or 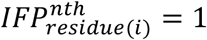.

### 4.3. MKI Database

Available MKIs were collected from review papers,^[15, 29]^ and scientific databases, including PubMed,^[57]^ and PDB.^[44]^ Databases were searched using the query keywords “macrocyclic”, “kinase”, and “inhibitors”. The search results were manually checked and all MKIs with nanomolar-level inhibition were collated, including SMILES format, primary target(s), assay data (*IC_50_*, *K_i_*, or *K_d_*), clinical states, PDB structures, and references. MKI scaffolds are obtained by manually cutting off the bridge linkers and then calculating the Rings_with_Linkers (RWL) scaffold using the tool strip-it.^[37]^ For every MKI and its corresponding RWL scaffold, the molecular properties were calculated using JChem^[58]^ and Mold2 for calculating Kier flexibility indices for molecular flexibility.^[59]^ The pairwise similarity of the RWL scaffolds was calculated using JChem with the molecular descriptor ECFP and Tanimoto distance. The RWL scaffolds were clustered using the average-linkage algorithm (**Table S4**) and shown using a circular dendrogram in RStudio (version 1.4.1106).

Based on the types and positions of atoms of the core structures interacting with the hinge, we sorted and clustered them into eight different types (**Table S8**). The Reingold-Tilford Tree network diagram was created using the networkD3 package in RStudio (version 1.4.1106) showing the 8 types of core structures. All compounds were illustrated with professional ChemDraw (version 20.0.0.38).

## 5 Supporting Information

**Table S1-S7** describes the MKI database. 95 nonredundant scaffolds, distributions of 10 clusters of scaffolds, physicochemical properties of MKIs, physicochemical properties of scaffolds, and physicochemical properties of MKIs and Ro5, respectively (xlsx). **Table S8** describes the 73 core structures (docx); **Figure S1** illustrates the RMSDs of all six MD systems for validating the MD processes (docx). The online MKI database is at https://zhengzhster.github.io/MKIs.

## 6. Acknowledgement

The authors acknowledge Research Computing at The University of Virginia for providing computational resources and technical support. This work described here was supported by the University of Virginia (P.E.B).

