## Supplementary material for "Rigid Scaffolds are Promising for Designing Macrocyclic Kinase Inhibitors": Figure S1 and Table S8

**Table S8. 73 Core Structures.**

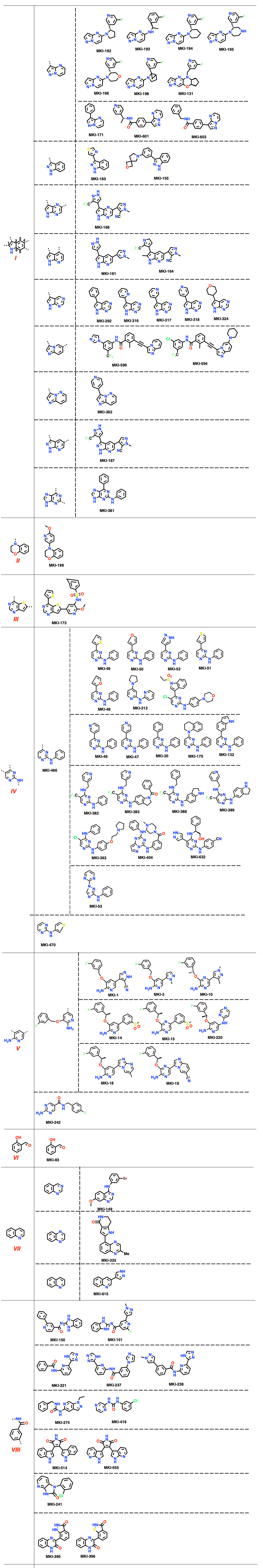

**Figure S1 continued**

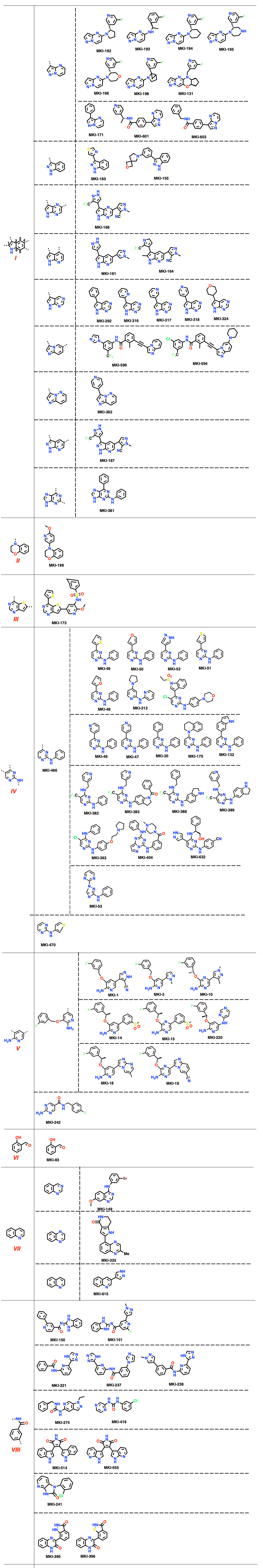

**Figure S1 continued**

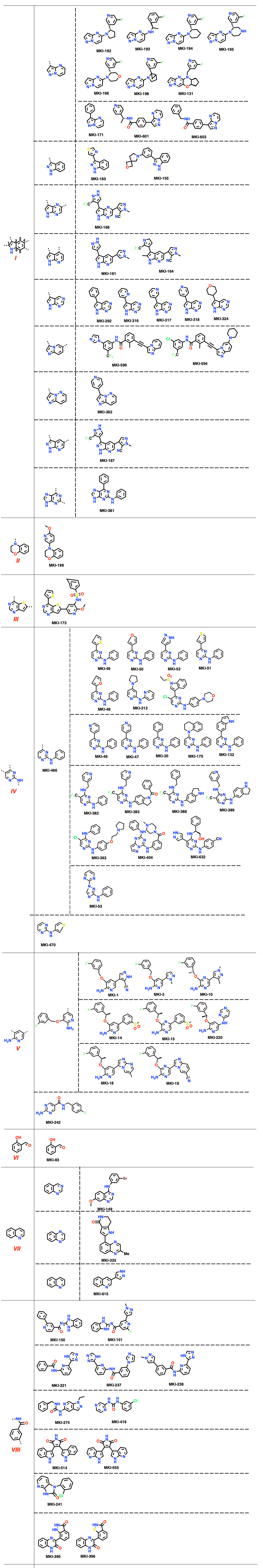

**Figure S1 continued**

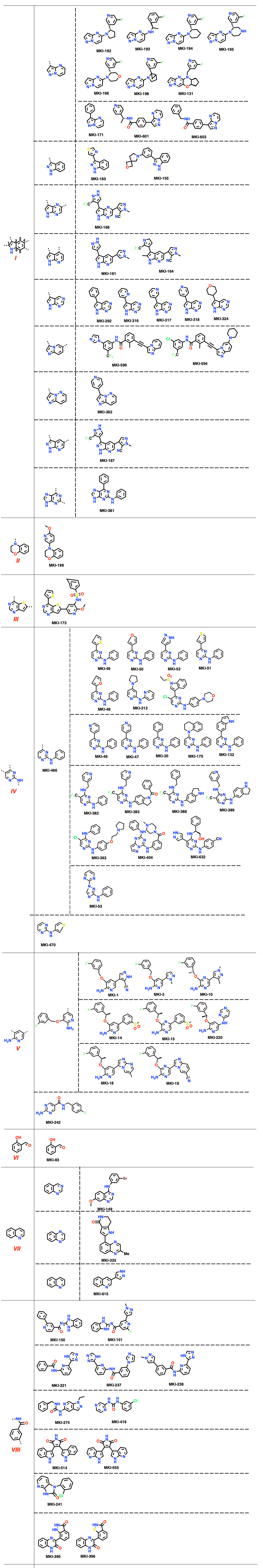

**Figure S1 continued**

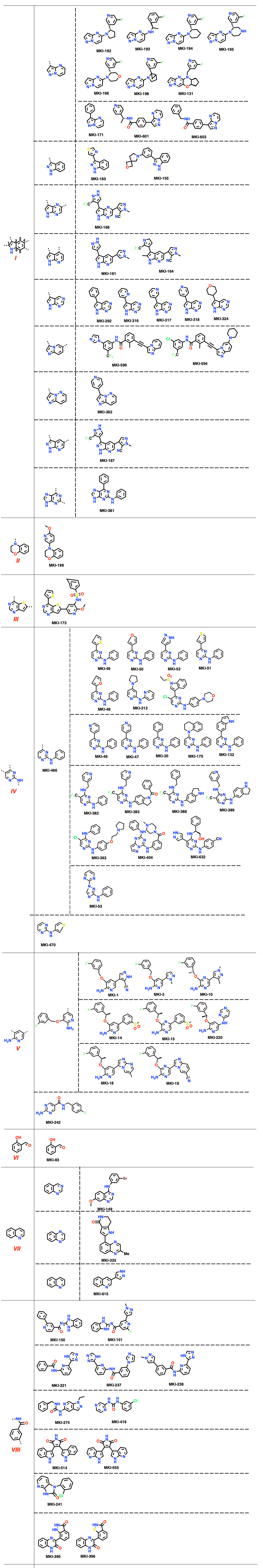

### **Figure S1. The RMSDs of six MD systems**

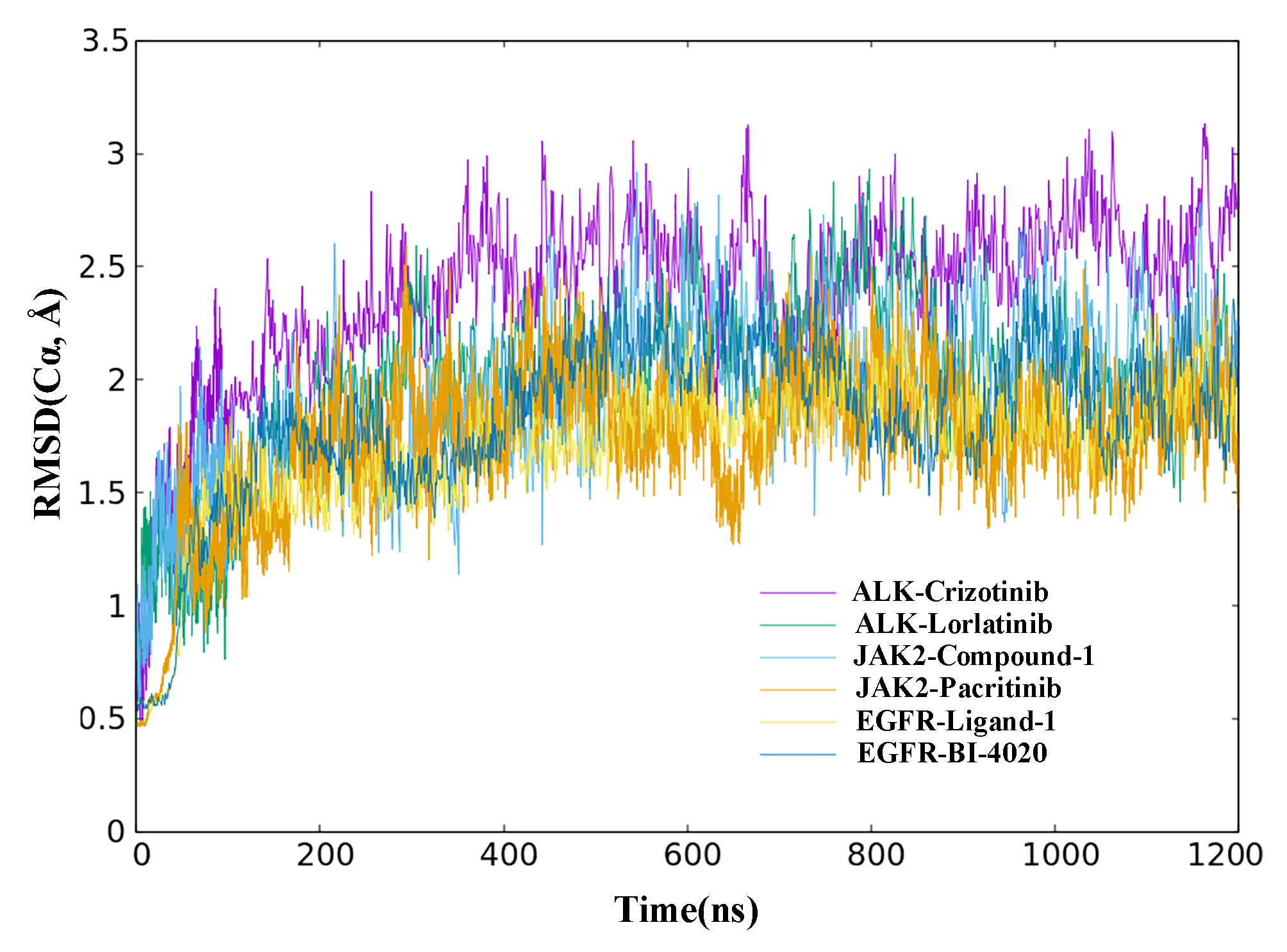
